# Human neutrophil response to Pseudomonas bacteriophages

**DOI:** 10.1101/786905

**Authors:** Dwayne R. Roach, Sylvie Chollet-Martin, Benoît Noël, Vanessa Granger, Laurent Debarbieux, Luc de Chaisemartin

## Abstract

The immune system offers several mechanisms of response to remove harmful microbes that invade the human body. As a first line of defense, neutrophils can remove pathogens by phagocytosis, inactivate them by the release of reactive oxygen species (ROS) or immobilize them by neutrophil extracellular traps (NETs). Although, recent studies have shown that bacteriophages (phages) make up a large portion of human microbiomes and are currently being explored as human antibacterial therapeutics, neutrophilic responses to phages are still elusive. Here, we show that exposure of isolated human resting neutrophils to high concentration of the *Pseudomonas* phage PAK_P1 led to a 2.8 fold increase in interleukin-8 (IL-8) secretion. Importantly, phage exposure did not further affect resting neutrophil apoptosis or induce necrosis, CD11 expression, oxidative burst, and NETs. Similarly, inflammatory stimuli activated neutrophil effector responses were unaffected by phage exposure. Our work suggest that phages are unlikely to inadvertently cause excessive neutrophil responses that could damage tissues and worsen disease. Because IL-8 functions as a chemoattractant directing immune cells to sites of infection and inflammation, phage-stimulated IL-8 production may boost host immune responses.

## 1. Introduction

As a key component in the innate immune system, neutrophils are generally considered as rapid responders in the first line of defense against pathogen invasion. They are the predominant population among leukocytes and can clear pathogens by a number of mechanisms, including phagocytosis, production of reactive oxygen species (ROS) and other antimicrobial products, and neutrophil extracellular traps (NETs) (1,2). The role of neutrophils has long been considered restricted to the initial phase of defense. However, recent evidence indicates that there is a functional heterogeneity and plasticity among neutrophils that shape both innate and adaptive immune responses by the secretion of cytokines (3,4). To this end, neutrophils are regularly associated with inflammation and disease, while also promoting inflammation resolution and homeostasis.

The human microbiota is the aggregate of more than 100 trillion symbiotic microorganisms that live on and within the body, including bacteria, archaea, eukaryotic viruses and bacteriophages (phages) (5,6). It is now acknowledged that the human microbiota affects host physiology to a great extent, including host immunity and homeostasis (5,7). Recently, resident phages – a diverse group of viruses that infect bacteria – community structure and composition have been shown to be altered during inflammatory diseases, such as inflammatory bowel disease (8,9), periodontal disease (10) and diabetes (11). Although phages are not human pathogens, Duerkop and Hooper hypothesize that phages may also trigger antiviral defenses (6) because they can elicit immune responses (12). More recently, phages were proposed to be implicated in the efficacy of fecal microbiota transplantations (13). This suggests phages have an unidentified role in affecting host immunity.

Moreover, phages has been proposed as a human antibacterial therapy, namely phage therapy, to address the rise of multidrug resistant infections (14,15). The mechanism by which phages exert their therapeutic action has generally been considered to be via their capacity to lyse bacterial cells. However, studies have shown that phages and the innate immune system work synergistically to eliminate bacterial infections (16,17). Interestingly, neutrophils were deemed necessary for the successful phage therapy in an animal model (17). Neutrophils may be keystone in determining the outcome of phage therapy; however, phage-neutrophil interactions are poorly studied.

Here, we describe the interactions between resting human peripheral blood neutrophils and the *Pseudomonas aeruginosa* infecting phage PAK_P1. We also determine the effect of phage exposure while activated neutrophils respond to several biological and chemical inflammatory stimuli.

## 2. Materials and Methods

### 2.1. Bacteria and phage culture

*P*. *aeruginosa* strain PAK was cultured in Luria Broth (LB; BD Biosciences) at 37°C. The pseudomonas phage strain PAK_P1 (17) was propagated on PAK grown in LB medium and purified following cesium chloride density gradient ultracentrifugation (18). After buffer (10mM Tris, 150mM NaCl, pH 7.5) dialysis, phages were passaged 3x through an endotoxin removal spin column (EndoTrap, Hyglos) to obtain residual endotoxin levels <1.5 EU mL^−1^ (EndoZyme II recombinant factor C, Hyglos). Phage stocks were 0.22 µm filter sterilized and enumerated on PAK. Before experimentation, phage numbers were adjusted in Hank’s balanced saline solution (HBSS no calcium, no magnesium; Fisher Scientific).

### 2.2. Neutrophil isolation and stimulation

Neutrophils were separated from human peripheral blood collected from healthy donors on EDTA by negative magnetic sorting (MACSxpress, Miltenyi Biotec) according to manufacturer’s instructions. Neutrophils were washed and resuspended at 3×10^6^ mL^−1^ in HBSS, and purity >98% and viability >99% was confirmed by flow cytometry. Equal volumes of phage solution or buffer control and neutrophils were mixed to give a final neutrophil concentration of 1.5×10^6^ mL^−1^. Inflammatory responses were triggered with either 25 nM phorbol myristate acetate (PMA), 5µM calcium ionophore (A23187), 20µM platelet activating factor (PAF), 100µg/mL Zymosan A, 5µg mL^−1^ Staphylococcus aureus peptidoglycan (PGN), or 100µg mL^−1^ ovalbumin/anti-ovalbumin immune complex at a 1:5 ratio in PBS. All stimuli were from Sigma-Aldrich.

### 2.3. Quantification of cell surface CD11b expression

Neutrophil CD11b expression was measured after 15 min saturation in 4% BSA PBS and membrane staining with PE-conjugated anti-CD11b (Becton-Dickinson) for 30 min at 4°C in darkness. Mean fluorescence intensity (MFI) of washed cells was measured on an Attune Nxt cytometer (ThermoFisher).

### 2.4. Phagocytosis quantification

Neutrophil phagocytosis was measured after 90 min incubation in darkness with increasing concentrations of pHrodo Red Zymosan BioParticles (ThermoFisher) conjugates at 5-50 µg mL^−1^, as per supplier’s instructions. Phagocytosis was monitored by an increase in particle fluorescence in acidic compartments using flow cytometry.

### 2.5. Oxidative burst response

ROS production was measured by chemiluminescence as previously described (19). Briefly, cells were incubated with phage with or without noxious stimuli for 40 min at 37°C in the presence of 100 mM luminol in a 96-well microplate (Corning). A TristarTM LB941 microplate reader (Berthold Technologies) measured luminescence every 1 min and area under the curve (AUC) was calculated for each sample tested in triplicate.

### 2.6. Neutrophil Extracellular Traps release

Extracellular DNA release was measured by Sytox Green fluorescence, as described previously (19). Briefly, the neutrophils were incubated with phage with or without inflammatory stimuli in presence of 5 µM of Sytox Green (Fisher Scientific) in a 96-well microplate (Corning) and samples were tested in triplicate. A microplate reader measured DNA release over 3 h and background fluorescence was subtracted from values.

### 2.7. Interleukin-8 (IL-8) production

Neutrophils were cultured for 18 h at 37°C in the presence of phages, alone or with PAF or PGN to stimulate the neutrophils. IL-8 levels were measured in cell-free culture supernatant by ELISA (hIL-8 Quantikine kit, Bio-techne), according to manufacturer’s instructions, and measured on a Multiskan EX spectrophotometer (Thermo Labsystems).

### 2.8. Apoptosis and necrosis

Neutrophils were incubated with phages for up to 18 h at 37°C. Cell apoptosis and necrosis were measured by annexin V/ 7-AAD (Annexin V Apoptosis Detection Kit I, BD Pharmingen), according to the manufacturer’s instructions. Cell viability was also evaluated using Trypan blue (Sigma-Aldrich) staining.

### 2.9. Statistical analyses

Experimental groups were analyzed by Mann and Whitney unpaired U-test and Kruskal-Wallis test. Results are shown as mean ±SEM. A p-value < 0.05 was considered statistically significant.

### 2.10. Ethics statements

Fresh whole blood samples were collected from healthy blood donors who gave their oral consent, in agreement with the French regulations and provided by Etablissement Français du Sang.

## 3. Results

### 3.1. Neutrophilic response to phages

We first established the characteristic neutrophil activities towards purified phage PAK_P1 compared to the HBSS diluent control. Phage PAK_P1 was found to not have a significant effect on neutrophil apoptosis after either short (3 h) or long (18 h) co-incubations (Fig. 1a & b). In addition, phages did not induce necrosis, even at the high cell:phage ratio of 1:10,000 (Fig. 1c and Suppl. Fig. 1). Phages also did not significantly induce neutrophils to upregulate CD11b cell surface expression (Fig. 1d), which regulates leukocyte adhesion and migration. Fig. 1e shows that phages did not induce ROS production, which is a crucial reaction that occurs in neutrophils to degrade internalized particles and microbes. Neutrophils have also been shown to kill pathogens outside of the cell, rather than engulfing them. This occurs by neutrophils releasing web-like structures of chromatin and granules, called NETs (1). Phage PAK_P1 alone did not trigger NETs (Fig. 1f). Together, this suggests human neutrophils were unresponsive to phage PAK_P1. However, we found that neutrophils secreted 2.8 fold higher amounts of IL-8 after 18 h of co-incubation with phage PAK_P1 when at the 1:10,000 cell:phage ratio compared to the control (Fig. 1g). Because IL-8 is involved in neutrophil activation and chemoattraction of other immune cells, human neutrophils may sense phage virions as foreign invaders.

**Figure 1:**
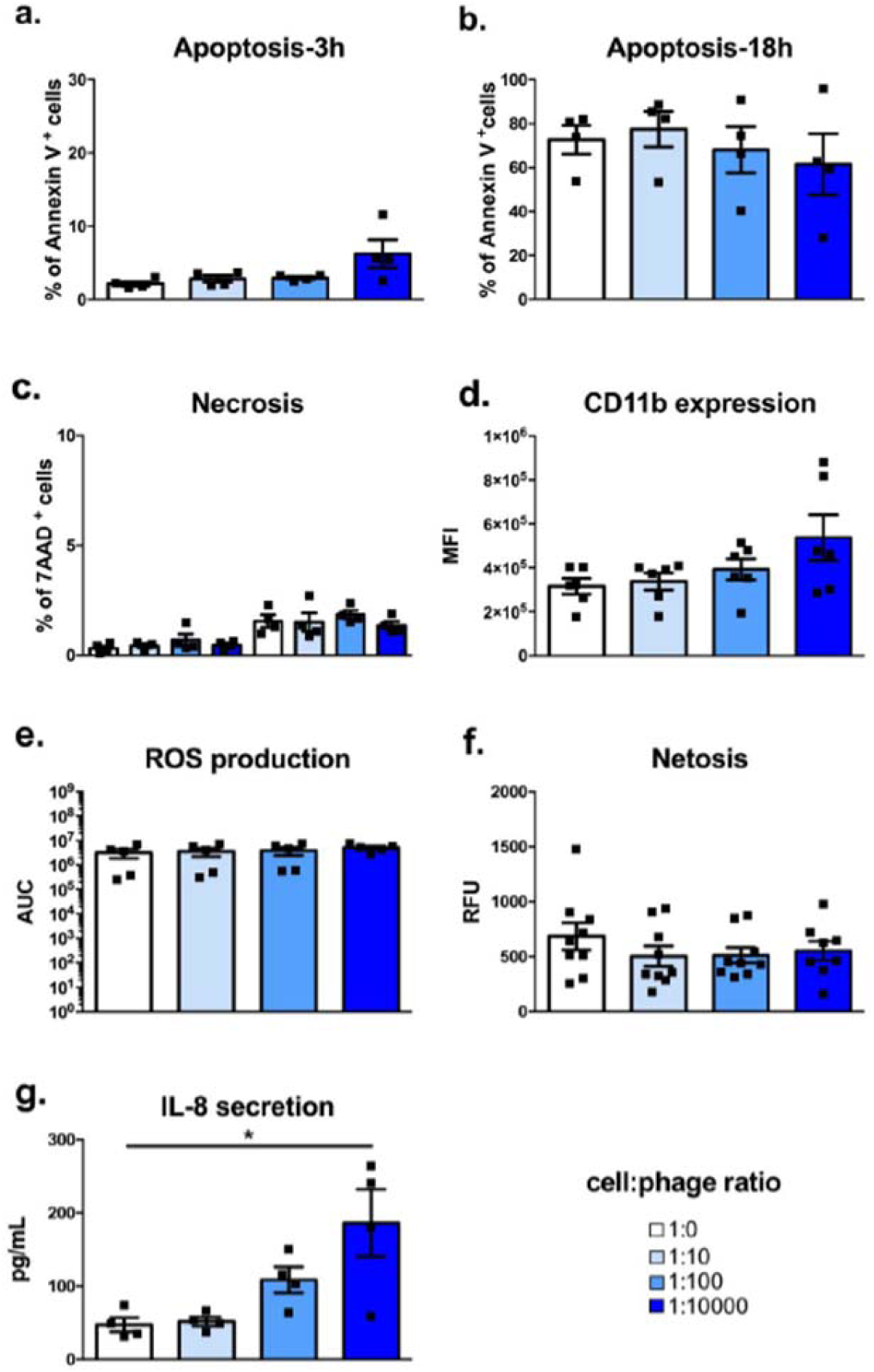
Resting human neutrophil responses to phage PAK_P1. Isolated human peripheral neutrophils were co-incubated with increasing amounts of purified phages. **(a&b)** Percent of annexin V positive neutrophils indicative of cell apoptosis after **(a)** 3h and **(b)** 18h co-incubation. **c)** Percent of 7-AAD positive neutrophils indicative of necrosis after co-incubation. **d)** CD11b expression as mean fluorescence intensity (MFI) after 18 h co-incubation. **e)** ROS production shown as luminescence area under curve (AUC) after 40 min co-incubation. **f)** Extracellular DNA release shown as relative fluorescence units (RFU) after 3 h co-incubation. **g)** Interleukin 8 (IL-8) secretion after 18 h co-incubation. Data shown as mean +SEM, n= 4-9 per group; *Kruskal-Wallis test p <0.05.

### 3.2. Neutrophilic responses to inflammatory stimuli are not influenced by phages

We then tested if phage exposure caused an overly lively neutrophil response towards inflammatory chemical (PMA and A23187), microbial (PGN and Zymosan), soluble immune complexes, and PAF stimuli. Phage PAK_P1 exposure did not affect the percent of pHRodo positive cells at either a low (5µg mL^−1^) or high (50µg mL^−1^) Zymosan coated particle dose (Fig. 2a), which suggests phagocytosis was not perturbed. Furthermore, phage exposure did not alter ROS production in response to Zymosan or PMA (Fig. 2b). Indeed, the combination of phages and an inflammatory stimulus could induce a more exuberant response like NETs (1). However, co-incubation did not change the amount of extracellular chromatin released from neutrophils in response to both strong (PMA and A23187) and weak (PAF and IC) stimuli (Fig. 2c and 2d, respectively). Interestingly, activated neutrophil IL-8 production in response to PAF and PGN did not increase with phage exposure (Fig. 2e), despite phages alone causing an increase in IL-8 production in resting neutrophils (Fig. 1g). Nevertheless, our results suggest that phage PAK_P1 exposure neither enhances nor dampens activated neutrophil responses.

**Figure 2:**
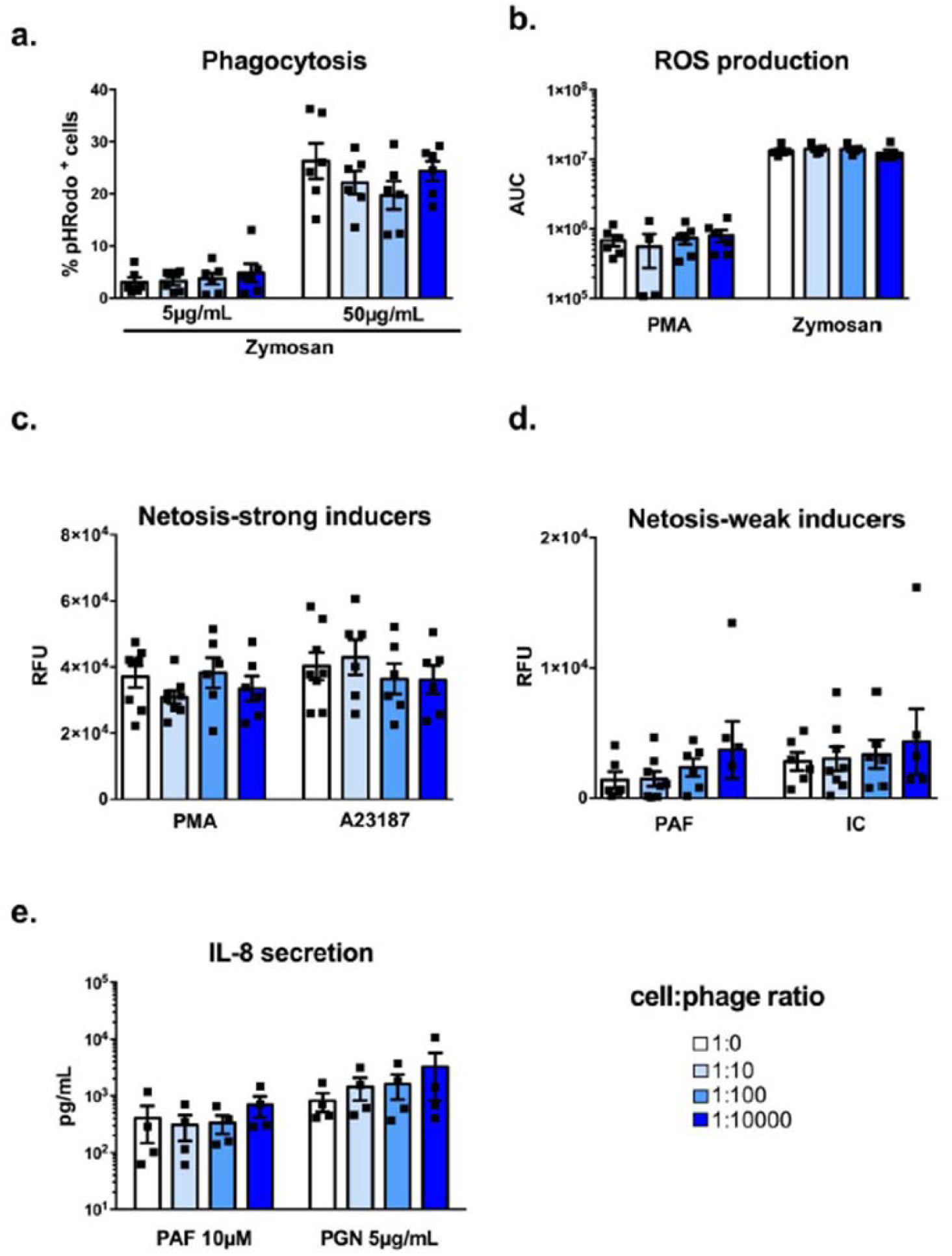
Activated human neutrophil responses to phage PAK_P1. Isolated human peripheral neutrophils were activated with either fungal glycan (Zymosan), phorbol myristate acetate (PMA), bacterial peptidoglycan (PGN), calcium ionophore (A23187), soluble immune complexes (IC) or platelet-activation factor (PAF) and co-incubated with increasing amounts of purified phages. **a)** Percent of pHRodo positive low (5 µg mL^−1^) or high (50 µg mL^−1^) Zymosan stimulated neutrophils after a 90 min co-incubation with phages. **b)** Mildly (PMA) or strongly (Zymosan) activated neutrophil ROS production shown as luminescence area under curve (AUC) after 40 min co-incubation with phages. **(c&d)** Strongly (PMA and A23187) **(c)** or weakly (PAF and IC) **(d)** activated neutrophil extracellular DNA release shown as relative fluorescence units (RFU) after 3 h co-incubation. **e)** Activated neutrophil interleukin 8 (IL-8) secretion after 18 h co-incubation. Data shown as mean +SEM, n= 4-8 per group.

## 4. Discussion

In this study, we show that a high concentration of purified *Pseudomonas* phage PAK_P1 can trigger low IL-8 production in freshly isolated human blood neutrophils. Nonetheless, phage exposure did not further affect resting neutrophil apoptosis or induce necrosis, CD11b expression, oxidative burst, and NETs. Similarly, activation-induced neutrophil effector responses were unaffected by phage exposure. Our findings agree with those from Borysowski *et al.* who similarly found that exposure to the *Staphylococcus aureus* phage A3/r did not increase granule marker expression in neutrophils (20). However, another study found that exposure to the E. coli phage T4, elicited a weak neutrophil oxidative burst (21). Furthermore, Miedzybrodzki *et al*. showed that co-exposure of the phage T4 with either *E. coli* or LPS derived from *E. coli*, had the beneficial effect of dampening neutrophil ROS production (22). Neutrophil antiviral defense remains controversial because it mediates both detrimental and beneficial effects to the host (23).

Human neutrophils generally cannot sense DNA viruses, largely because of lacking the DNA sensing Toll-like receptor 9 (23). DNA phages are the most abundant viruses in microbiomes and are used as therapeutic agents; therefore, it is unlikely neutrophils sense phages by their DNA content. Despite that, neutrophils have been implicated in the anti DNA-viral immune response (23). For example, herpes simplex virus (HSV) can cause a rapid neutrophil infiltration to the site of infection, and neutrophil depletion results in increased HSV load (24). Epstein-Barr virus (EBV) induces IL-8 production in neutrophils, which was found to be dependent on the interaction between the viral envelope and cell surface (25). A major difference however is that DNA phages lack a viral envelope. Another key difference is that HSV and EBV elicits neutrophilic effector responses (e.g., apoptosis and oxidative burst) in parallel to IL-8 production (24,25). This was not observed during phage PAK_P1 exposure.

A caveat of deciphering anti-phage immune response is that phages propagate in bacteria. Importantly, neutrophils are highly bacterial responsive phagocytes, expressing an abundance of bacteria-specific receptors, including the endotoxin (i.e. LPS) sensing Toll-like receptor 4 (1). Thus, trace amounts of LPS in the phage preparation might be responsible for IL-8 secretion in neutrophils (26). However, LPS tends to enhance neutrophil sensitivity to subsequent inflammatory stimuli (27), which was not observed after exposure to the phage PAK_P1 preparation. It has also been shown that phage tail fibers can bind free LPS making it less inflammatory (28), which could thereby scavenge trace endotoxin after the LPS removal procedures used in this study during phage purification.

Although neutrophils are the first and predominant immune cell population recruited to an affected site after infection, it remains unclear if this occurs when triggered by phage signals. Importantly, our results suggest that phages are unlikely to stimulate neutrophils to inadvertently release their intracellular toxic contents and cause collateral tissues damage. As a major component of the human microbiota (6), phages do not appear to play a major role in diseases where neutrophils have been implicated (29,30). As antibacterial therapeutic agents, phages have been shown to work in concert with neutrophils to cure bacterial infection (17). But, phages do not appear to enhance neutrophil effector responses during treatment. However, because IL-8 has other biological functions besides a central role in inflammation, such as immune cell chemotaxis, angiogenesis, and hematopoiesis (4), phage-stimulated IL-8 production may boost immune responses.

The interaction between human immune cells and phages that either make up part of the human microbiota or are administered as human therapeutics, remains underappreciated. A better understanding of phage-stimulated immune responses could open new treatment options for infectious and inflammatory diseases.

## Author contributions

DRR, SCM, LDC and LD jointly conceived the experiments. DRR, BM, and LDC performed experiments. DRR, SCM, LDC and LD analyzed the data and prepared the manuscript.

---

**Supplementary Figure 1:**
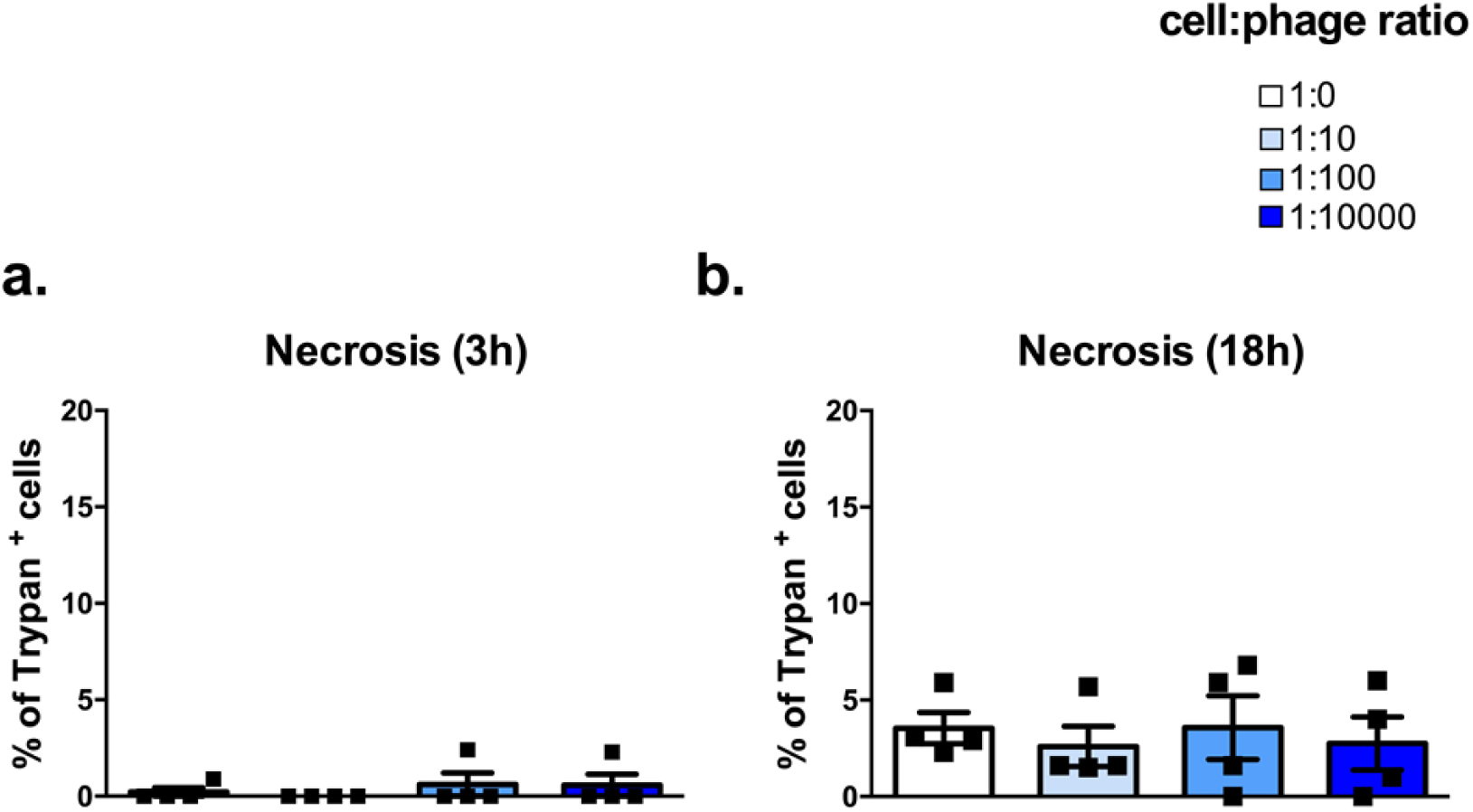
Resting human neutrophil necrosis after phage PAK_P1 co-incubation. Percent of Trypan blue positive human peripheral neutrophils after **(a)** 3 h and **(b)** 18 h co-incubation with increasing amounts of purified phages. Data shown as mean +SEM, n= 4 per group.

